# The distribution of waiting distances in ancestral recombination graphs and its applications

**DOI:** 10.1101/2020.12.24.424361

**Authors:** Yun Deng, Yun S. Song, Rasmus Nielsen

## Abstract

The *ancestral recombination graph* (ARG) contains the full genealogical information of the sample, and many population genetic inference problems can be solved using inferred or sampled ARGs. In particular, the waiting distance between tree changes along the genome can be used to make inference about the distribution and evolution of recombination rates. To this end, we here derive an analytic expression for the distribution of waiting distances between tree changes under the sequentially Markovian coalescent model and obtain an accurate approximation to the distribution of waiting distances for topology changes. We use these results to show that some of the recently proposed methods for inferring sequences of trees along the genome provide strongly biased distributions of waiting distances. In addition, we provide a correction to an undercounting problem facing all available ARG inference methods, thereby facilitating the use of ARG inference methods to estimate temporal changes in the recombination rate.

## 1 Introduction

At each position of the genome, the relationship among individuals in a sample can be characterized by a tree, known as a genealogical or coalescent tree, and it can be regarded as the result of a generative process called the *coalescent* (Kingman, 1982a,b). In the presence of recombination, the genealogies can vary at different positions of the genome, and the *ancestral recombination graph* (ARG) — the structure which fully describes the joint distribution of coalescent trees along the genome — provides all the information about the genealogical history of a sample, including the locations of recombination events. The full ARG can be also seen as the result of generative process, *the coalescent with recombination* (Griffiths, 1981; Hudson, 1983). Although it is straightforward to simulate under this process (Hudson, 2002; Kelleher et al., 2016), inferring ARGs from population genetic variation data is a very challenging problem. Indeed, much algorithmic work has been done on reconstructing parsimonious histories with recombination (Hein, 1993; Gusfield et al., 2003; Song and Hein, 2003; Wang et al., 2001; Lyngsø et al., 2005; Gusfield, 2014; Ignatieva et al., 2020) and sampling ARGs under the coalescent with recombination (Griffiths and Marjoram, 1996; Fearnhead and Donnelly, 2001; Jenkins and Griffiths, 2011; Rasmussen et al., 2014).

The coalescent with recombination (Griffiths, 1981; Hudson, 1983) was originally formulated as a stochastic process over time, but Wiuf and Hein (1999) later showed that it can also be reformulated as a spatial process along the genome. This spatial process is not Markovian, because of the long-range dependency caused by coalescence events between distant genomic positions. However, constraining the spatial process to be Markovian (McVean and Cardin, 2005; Marjoram and Wall, 2006; Hobolth and Jensen, 2014) has led to useful, practical approximations of the full coalescent with recombination while retaining accuracy in many aspects. The first Markovian approximation is called the *sequentially Markovian coalescent* (SMC) (McVean and Cardin, 2005), and a subsequent improvement (Marjoram and Wall, 2006), known as SMC’, incorporates an additional class of genealogical events.

The Markovian approximations have successfully been applied in the estimation of changes in population size (e.g., Li and Durbin (2011); Schiffels and Durbin (2014); Terhorst et al. (2017)), by representing the genealogy as states of a Hidden Markov Model (HMM), whose transition probabilities then can be calculated under the SMC or SMC’ assumptions. However, these methods are restrained to at most analyzing a few individuals due to the exploding size of the state space with increasing sample size. However, recently several different methods for inferring genealogies in models with recombination have been proposed (e.g., Rasmussen et al. (2014); Kelleher et al. (2019); Speidel et al. (2019)). *ARGweaver* by Rasmussen et al. (2014) is capable of full posterior sampling of ARGs under the SMC or SMC’ approximations using Markov Chain Monte Carlo (MCMC), but becomes prohibitively slow for large sample sizes (typically > 50). *Relate* by Speidel et al. (2019) and *tsinfer* by Kelleher et al. (2019) are capable of inferences for larger sample sizes, but do not perform full posterior sampling. Both methods only provide a single estimate of the tree topology, although *Relate* also samples coalescent times using MCMC, conditionally on the estimated local coalescent tree topology.

### 1.1 Motivation

In the SMC, the SMC’, and the full coalescent process with recombination, the waiting distance *d* until the next recombination event along the chromosome is exponentially distributed, conditionally on the total tree length 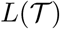 of the current tree 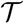:

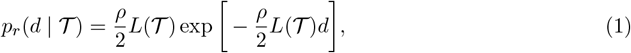

where 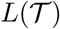 is in coalescent unit and 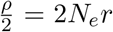 denotes the population-scaled recombination rate per bp. SMC’ provides a closer approximation to the full coalescent with recombination than does SMC, as the former allows for events in which a lineage splits off of and coalesces back to the same branch (type 1 in Figure 1a). This type of event occurs with particularly high probability when the sample size is small (Wilton et al., 2015). However, it does not change the genealogy (second tree in Figure 1b) and is therefore not sampled in *ARGweaver* or reported in the output of *msprime* simulations (Kelleher et al., 2016). Therefore, the waiting distance between adjacent trees in *ARGweaver* or *msprime* will not follow the exponential distribution shown in (1) (Figure 2a). Furthermore, there are two additional types of events (types 2 and 3 in Figure 1a) that may occur in both SMC and SMC’ approximations where some coalescence times change, but not the tree topology (Hudson and Kaplan (1985) considered different recombination types along this line and studied their statistical properties, and Hein et al. (2004) refined this classification). Hence, the waiting distance distribution until next topology change is even further biased away from (1) (Figure 2c), and similar problem was explored in the context of incompatibility by Hudson and Kaplan (1985).

**Figure 1:**
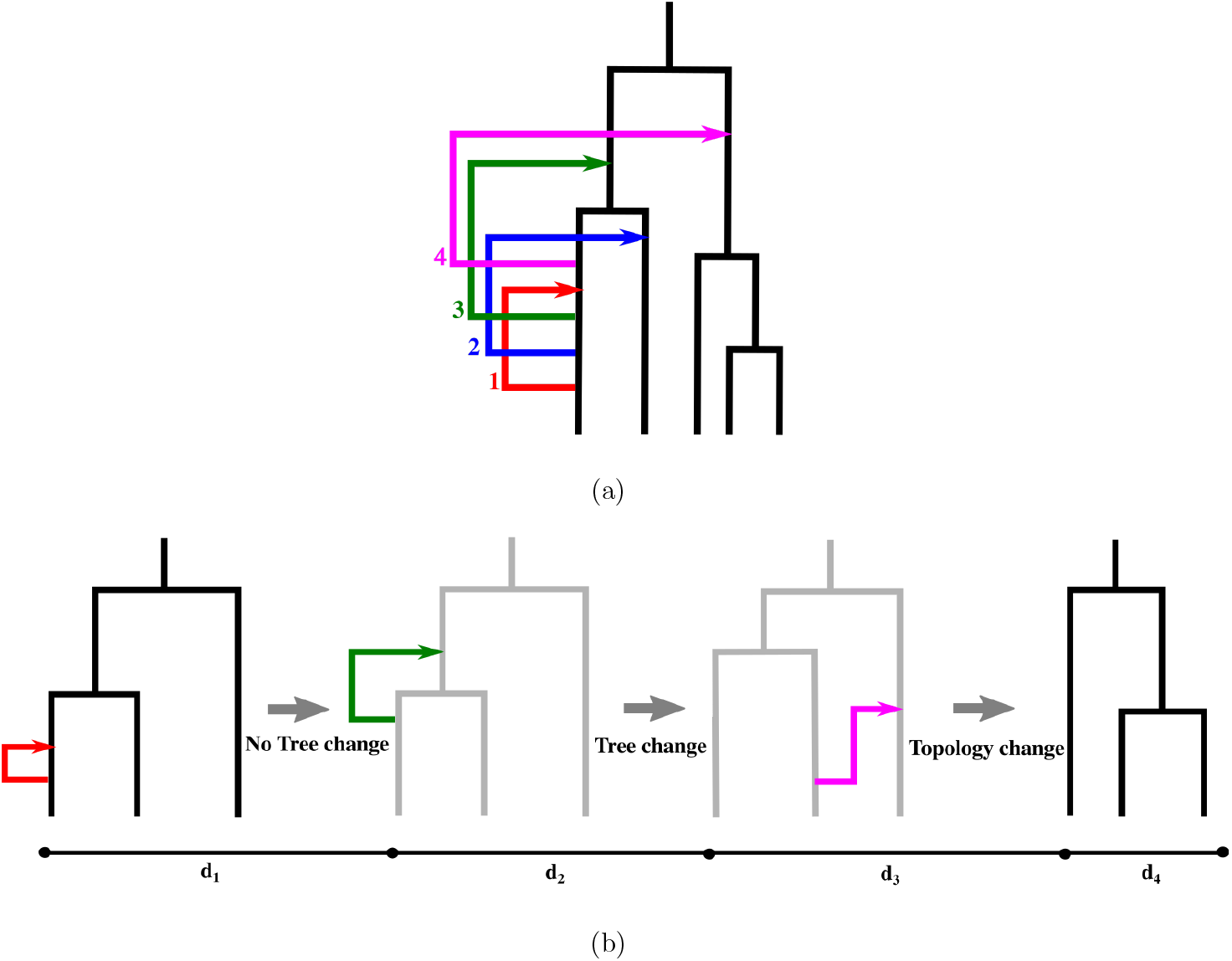
Difficulties in genealogy inference. (a) Different types of recombination events, in which type 1 does not change the tree, type 2 and 3 change the tree but not the topology, and type 4 changes the topology; (b) Illustration of tree transition omission in tree inference methods, in which the shaded trees are harder to detect as they are not produced by topology changes.

**Figure 2:**
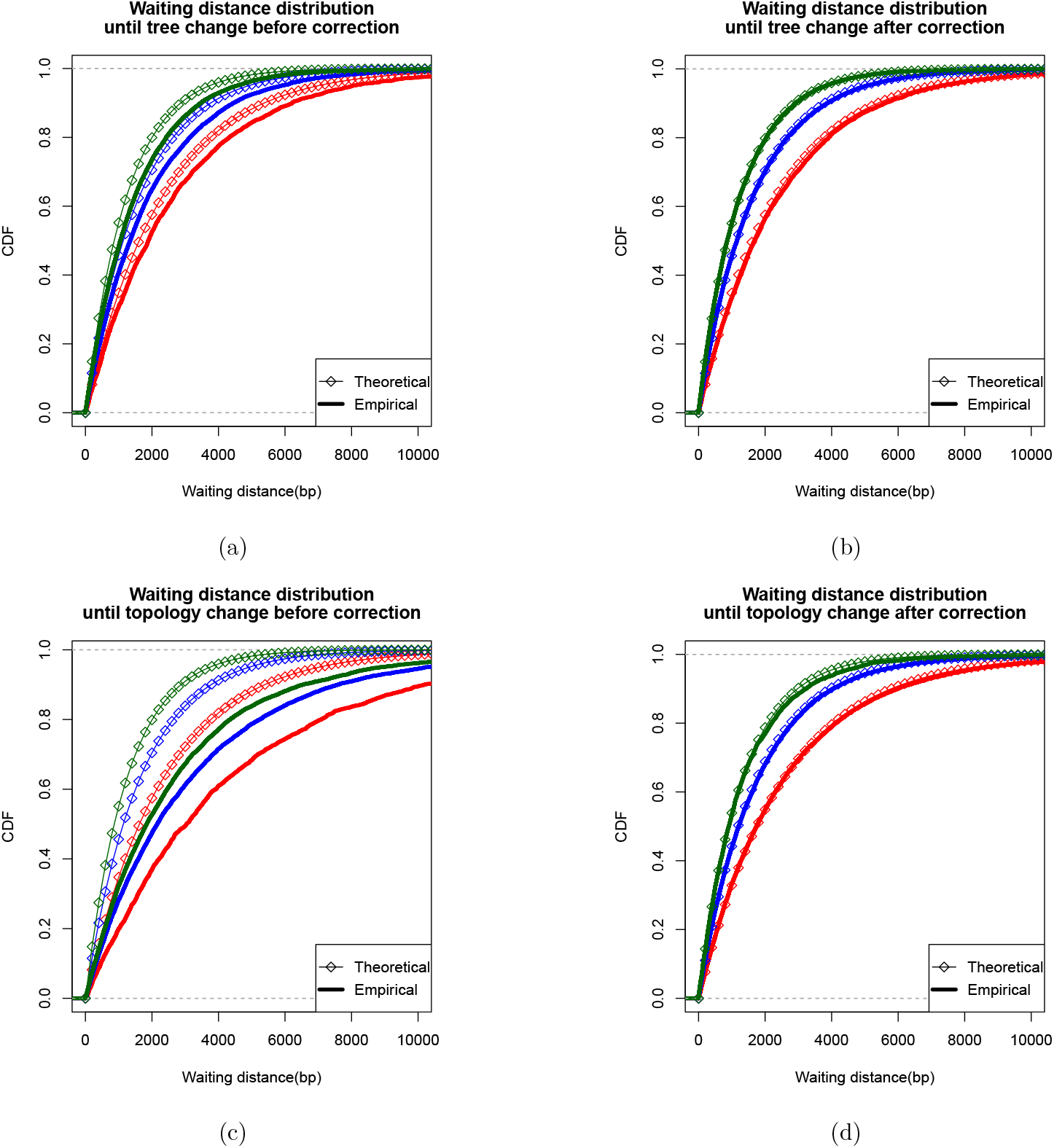
The waiting distance distribution until tree or topology changes. Simulations are done in *msprime* using *n* = 8,*N*_e_ = 1 × 10^4^,*μ* = *r* = 1 × 10^-8^ and the distribution of simulated waiting distances until tree or topology changes (solid lines) are compared to (1) and (4), (5) (squared lines). [Green, blue, and red mean the conditioned quantity to be within (3 × 10^4^, 5 × 10^4^), (5 × 10^4^, 7 × 10^4^), (7 × 10^4^,9 × 10^4^) respectively.] (a) The waiting distance until tree change differs from the simple exponential distribution in (1) (conditioning on 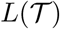); (b) The waiting distance until tree change matches the corrected exponential distribution of (4) (conditioning on 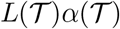); (c) The waiting distance distribution until topology change is quite different from the simple exponential distribution in (1) (conditioning on 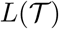); (d) The waiting distance distribution until topology change matches the corrected exponential distribution of (5) (conditioning on 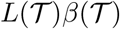).

**Figure 3:**
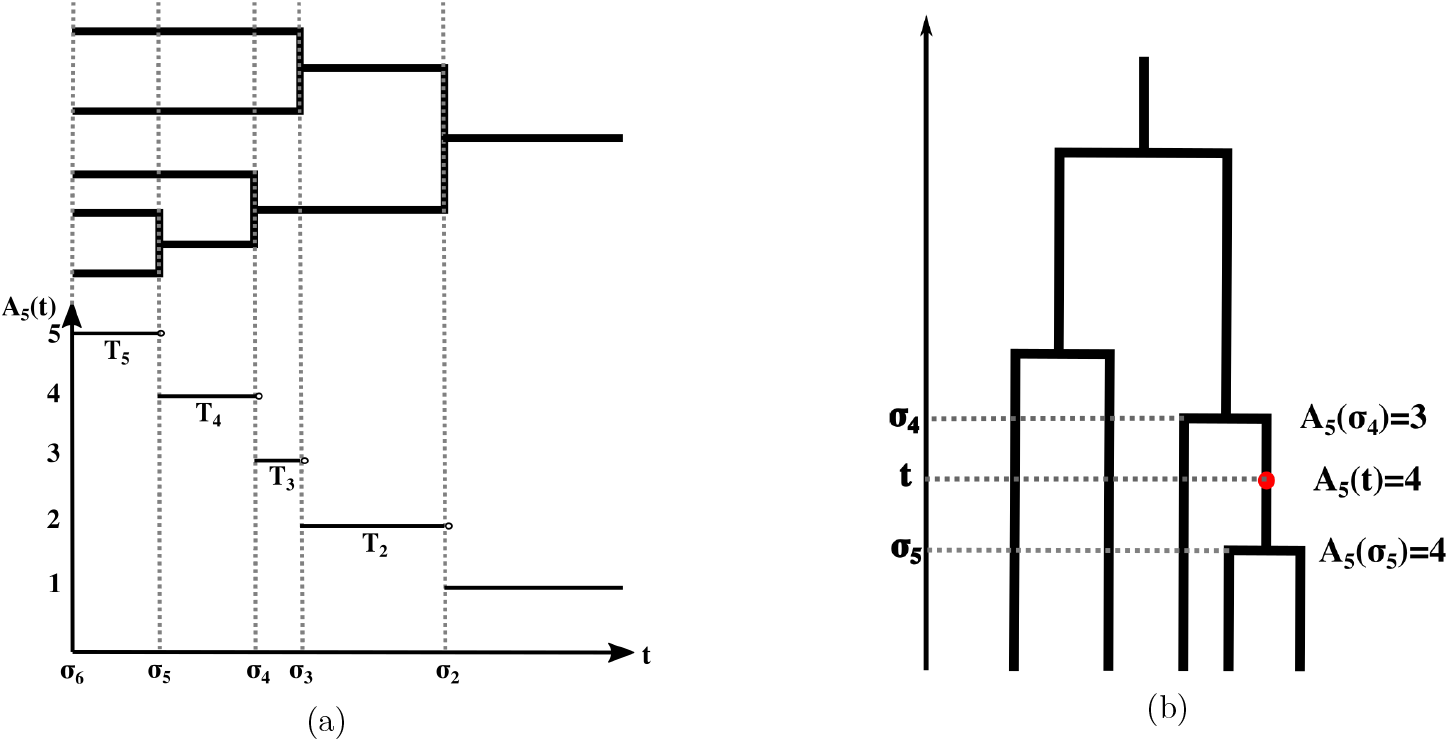
Illutration of our notational convention. (a) *A_n_*(*t*), *T_i_* and *σ_j_* are illustrated for an example genealogy with sample size *n* = 5. (b) A recombination event occurs on a branch at time *t*, and the corresponding upper and lower times of the branch are 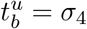 and 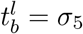.

To use the waiting distance distribution between trees inferred by SMC or SMC’ models, or as reported by common simulation programs such as *msprime*, for understanding patterns of recombination, or for validating the accuracy of inference methods or simulation methods, it is necessary to understand the distribution of waiting distances between tree topology changes induced by events of type 1, 2, or 3 in Figure 1a.

In this paper we derive the waiting distance distributions for the SMC’ model and we use these results to benchmark three common methods for inferring tree topologies along the length of a chromosome: *ARGweaver* (Rasmussen et al., 2014), *Relate* (Speidel et al., 2019), and *tsinfer* (Kelleher et al., 2019). We also illustrate the utility of the results, when used in combination with methods for inferring trees, for inferences regarding temporal changes in recombination rate.

## 2 Theoretical Results

### 2.1 Notation

Suppose we have a sample of size *n* from a diploid population with constant effective population size *N*_e_ (all times mentioned henceforth will be in coalescent units of 2*N*_e_ generations). We use *T_i_* to denote the length of the epoch when there are *i* ancestral lineages in the coalescent tree and introduce the notation

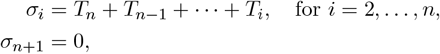

to denote the times when coalescence occurs. *A_n_*(*t*) will denote the number of lineages at time *t* ancestral to the sample. There are 2*n* – 2 branches in the coalescent tree before the most recent common ancestor (TMRCA), and we index them arbitrarily by *b* = 1,…, 2*n* – 2. For each branch *b*, we use 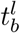 and 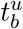 to denote its lower and upper times, respectively.

### 2.2 Waiting distance distribution until next tree change

The waiting distance until next tree change can be modeled as a waiting distance in a thinned Poisson process, where we color the events differently depending on whether they produce identical trees. Since the intensity of the un-thinned process is just the product of the tree length and the recombination rate, the only thing which needs to be identified is the thinning parameter, the probability of a recombination leading to a tree change. A simpler version of the problem was previously solved using the same idea for the waiting distance distribution until a TMRCA change for *n* = 2 (Carmi et al., 2014).

Now suppose the current tree is 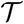, and the recombination happened on branch *b* at time *t*, we have the following result regarding the probability of this recombination not changing the tree under the SMC’ model (a type 1 event):

#### Proposition 1.

*Under the SMC’, the probability of a recombination not to change the tree, given its breakpoint on branch b at time *t* on tree 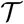 is*:

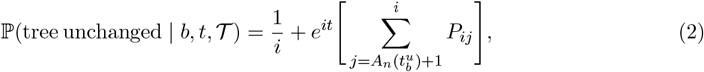

where

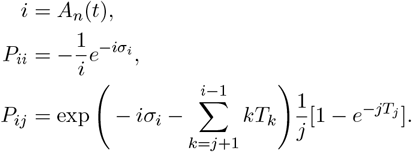

By a change in tree, we here mean a change in at least one coalescence time, but the topology may stay the same. This result, which is proved in the Appendix, shows that the probability of a recombination event being detected depends on where it occurs in the tree. We will also later show that this result facilitates inferences of temporal changes in recombination rates using inferred ARGs. We also note that under the SMC model 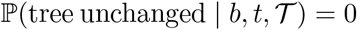.

The unconditional probability of type 1 event on a branch *b* under the SMC’ can then be found by integrating over the breakpoint time, *t*, with respect to its conditionally uniform distribution on the branch:

#### Proposition 2.

*Under the SMC’, the probability of a recombination happening on branch b not to change the tree is*:

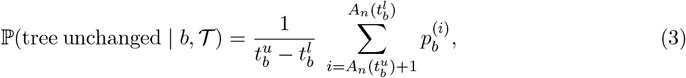

where

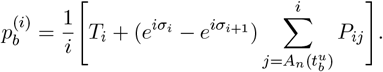

Now, the probability of a recombination event not changing the tree can simply be given by a weighted sum of the probabilities in (3) over all branches in the tree, weighting by the branch lengths:

#### Theorem 1.

***(tree-unchanging probability)** Under the SMC’, the probability of a recombination event not changing the tree is*:

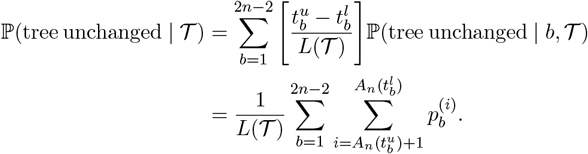

The waiting distance between tree changes under the SMC’ is then determined as follows.

#### Theorem 2.

***(waiting distance distribution until next tree change)** Under the SMC’, the distribution of waiting distance until next tree change given the current tree 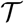 is given by*:

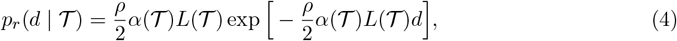

where

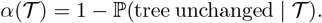

Note that the waiting distance until next tree change is still exponential, but the intensity is reduced by a factor of 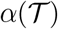. In Figure 2b we show that this expression indeed accurately describes the waiting distance distribution observed in *msprime* simulations.

### 2.3 Waiting distance distribution until next topology change

Although inferring every tree change would be desirable for ARG inference, it is notoriously hard, especially when scaling to hundreds, or thousands of genomes. Two of the most recent genealogy inference programs, *Relate* and *tsinfer*, use efficient approximations to infer genomic series of trees. However, inferences of tree changes are mostly guided by information regarding topologies, and neither method is designed to detect changes in the tree that does not involve a change in topology. For benchmarking and comparing these, and other programs, is therefore also important to understand the distribution of waiting distances between changes in topology.

To derive the waiting distance distribution between topology changes, we will again use the idea of a thinned process. The quantity of interest is the probability that a recombination event will change the tree topology. This probability can be calculated in very similar manner to that in Theorem 2, except that there are two more types of events of recombination and coalescent to consider (type 2 and 3) so some extra bookkeeping is needed. In Appendix A.2, we prove the following result:

#### Theorem 3.

***(topology-unchanging probability)** For a given branch b in tree 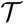, let b’ denote the branch that b coalesces with and let c denote their parental branch. Then, under the SMC’, the conditional probability that a recombination event will not change the tree topology given the current tree 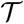 is*

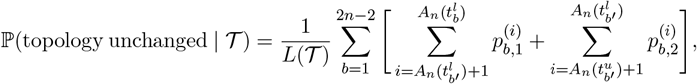

where

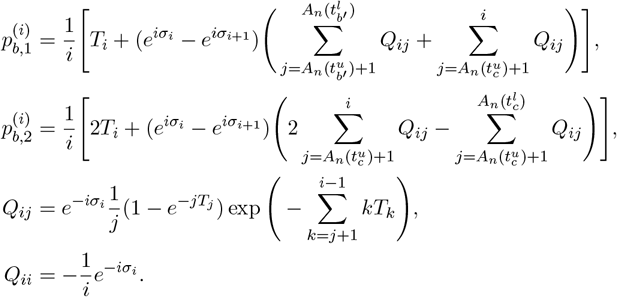

Unlike the result for the distribution of waiting distance until next tree change, the waiting distance to the next topology changing event is not exponentially distributed because of the possibility of recombination events that change the tree but do not change the topology. Such events will change the intensity of the process so that it is no longer time-homogenous. Arriving at an exact formula is, therefore, difficult. However, we notice that there is a rather accurate approximation method based on the following empirical observation:

#### Concentration phenomenon

The product of total tree length and the topology-changing probability remains approximately the same after a topology-unchanging recombination.

Notice in Figure 4 that while both the tree length, 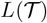, and the probability of a topology changing recombination event, 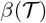, after a topology-unchanging event, has relatively high variance, the variance of their product is much smaller. In other words, a recombination+coalescent event that increases the total tree length will decrease the probability that the next event is topology changing and *vice versa*. The net effect is that the product of the two, 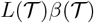, is much more stable than either are individually. Using this concentration relation we can build an approximation for the distribution of waiting distances until next topology change:

**Figure 4:**
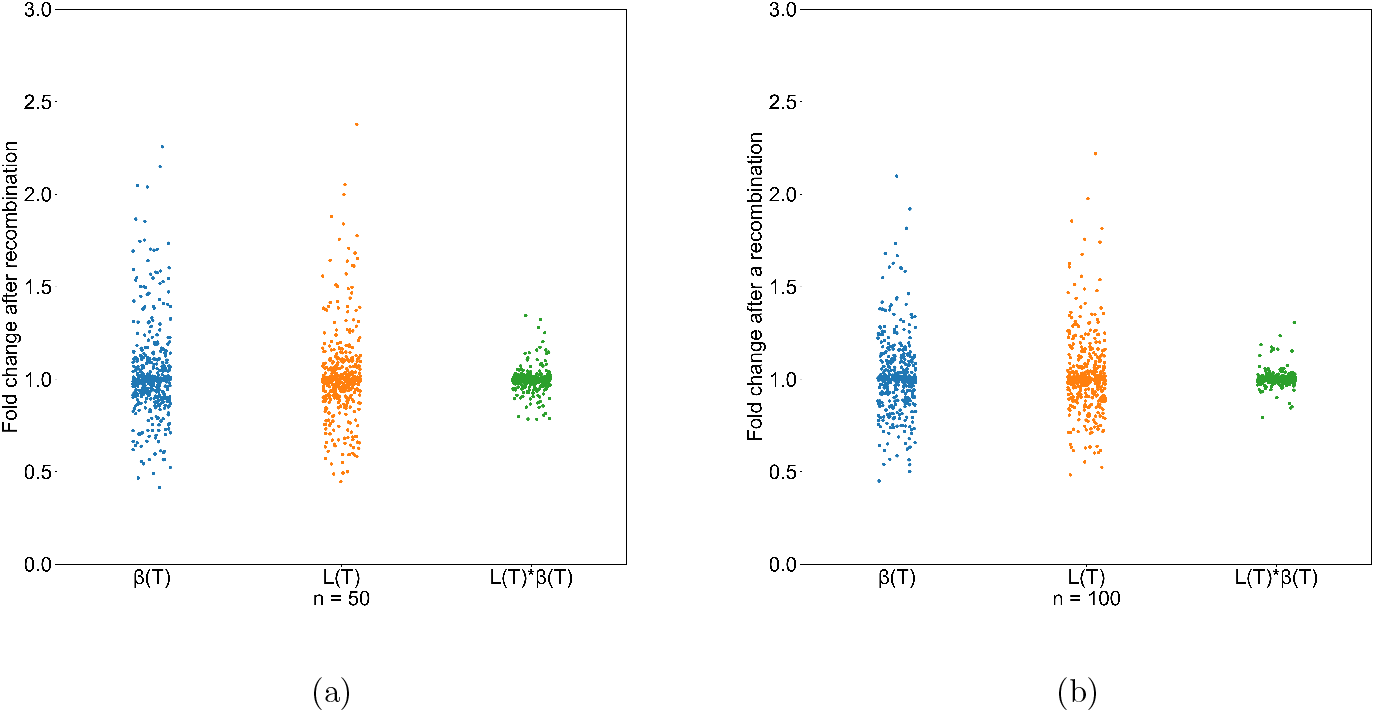
Concentration of 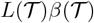 after a topology-unchanging recombination. Simulations are done using *msprime* under *n* = 20 and *n* = 50, in which the fold change of 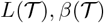 and 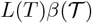 after recombination-unchanging recombinations are calculated. Although neither 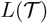 nor 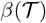 remains stable after a topology-unchanging recombination, their product remains approximately same with high probability. This relation is shown here rather empirically with simulation results using sample size *n* = 20 (a) and *n* = 50 (b).

#### Observation

**(Waiting distance distribution until next topology change)** The distribution of waiting distance until next topology change can be approximated by:

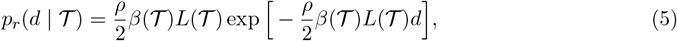

where

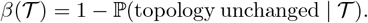

## 3 Applications

### 3.1 Benchmark of tree inference methods

One application of the theory introduced in this article is the use for benchmarking of some ARG/-genealogy inference methods. If the method detects every tree/topology change, or if it samples from a correct posterior distribution of waiting distances, then the waiting distance between adjacent trees should be described by (4) or (5).

To investigate this, we did simulations with realistic choices of parameters (*μ* = *r* = 1 × 10^-8^, *N*_e_ = 1 × 10^4^ with 8 haplotypes) and used *ARGweaver, Relate*, and *tsinfer* to infer the genealogy. Then we compared the distribution of waiting distance between adjacent trees in the output with our theoretical prediction (4) or (5).

Although *ARGweaver* does not report some tree transitions caused by type 1 recombination (Figure 5a), causing the waiting distance distribution to be different from the exponential distribution in (1), it is indeed capable of doing approximately posterior sampling of tree changes following (4) (Figure 5d). However, we observe a very small bias which might be due to the discretization in *ARGweaver*.

**Figure 5:**
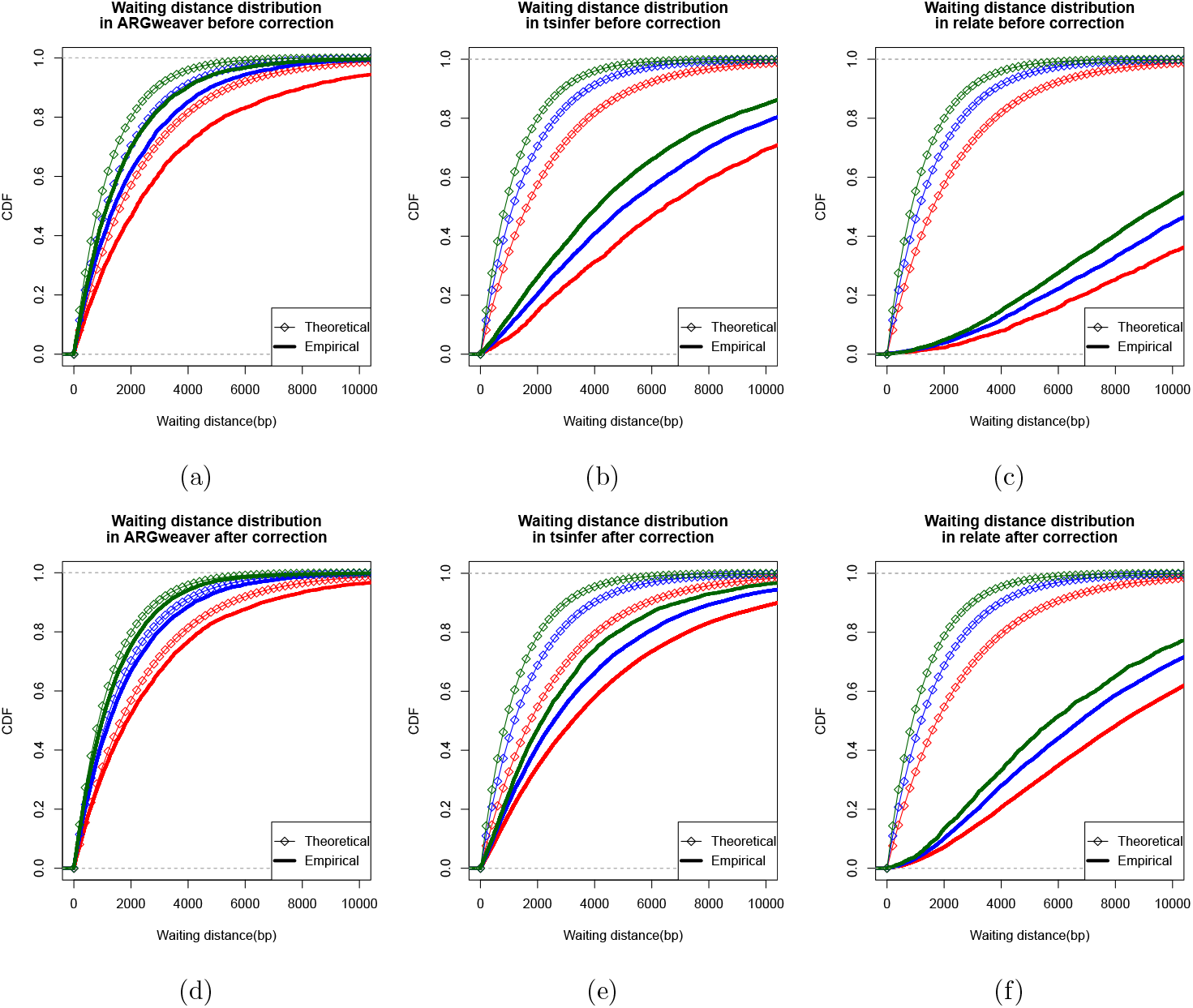
The waiting distance distribution in *ARGweaver, tsinfer* and *Relate*. Simulations are done in *msprime* using *n* = 8,*N*_e_ = 1 × 10^4^,*μ* = *r* = 1 × 10^-8^ and the distribution of waiting distances between adjacent trees in the outputs (solid lines) are compared to (1) and (4)/(5) (squared lines). Green, blue, and red mean the conditioned quantity to be within (3 × 10^4^, 5 × 10^4^), (5 × 10^4^, 7 × 10^4^), (7 × 10^4^, 9 × 10^4^). The conditioned quantities are 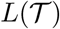 in (a), (b) and (c), 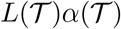 in (d), and 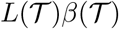 in (e) and (f). (a) The waiting distance distribution in *ARGweaver* is not well-characterized by the simple exponential distribution (1); (b) The waiting distance distribution in *tsinfer* is biasing away from the exponential distribution even more; (c) The waiting distance distribution in *Relate* is significantly biasing away from the exponential distribution; (d) The correction (4) provides a much better fit to the waiting distance distribution in *ARGweaver*; (e) The correction (5) is closer to the empirical distribution in *tsinfer*, but the correction doesn’t completely solve the problem; (f) The correction (5) on *Relate* helps little.

However, both *Relate* and *tsinfer* are undersampling trees. Under realistic choices of mutation and recombination rate of *μ* = *r* = 1 × 10^-8^ with 8 haplotypes, the waiting distance distribution in *tsinfer* is quite different from (5), suggesting that *tsinfer* undersamples toplogy changes and overestimates waiting distances (Figure 5e). *Relate* has an even stronger tendency to undersample topology changes and to overestimate waiting distances (Figure 5f).

### 3.2 Inference of temporal variation of recombination rate

It is generally hard to study the evolution of recombination rate through time without specific reference to ARGs. After sampling the ARG of a region using a program that accurately samples tree changes, such as *ARGweaver*, a naive way of estimating time-specific recombination rates is to count the number of recombination happening within each time interval, divided by the total branch length appearing in the interval. In order to test this procedure, we simulated data under a mutation rate of *μ* = 1 × 10^-8^ and a constant recombination rate *r* = 1 × 10^-8^ with 8 haplotypes of 1Mb. However, the naive estimation procedure provides biased estimates in at least two aspects (Figure 6): first, the inferred rates are significantly lower than the true values; second, the inferred trajectory is not constant, leading to a false conclusion of temporal changes in the recombination rate.

**Figure 6:**
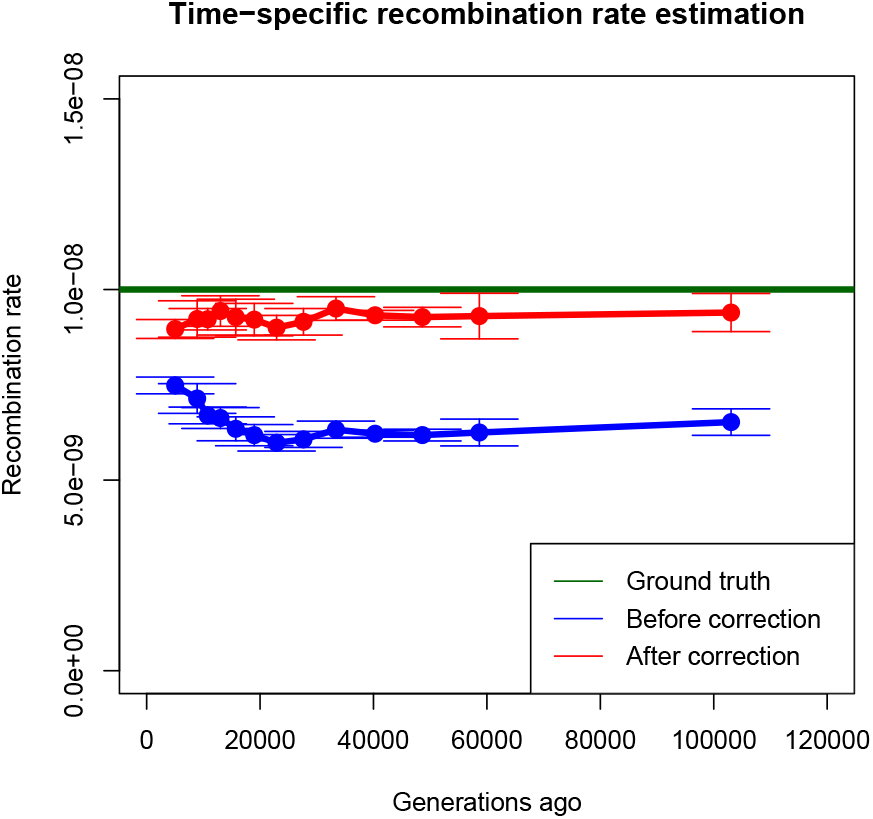
The time specific recombination estimation before and after correction using (2). The naive estimation without correction (blue line) is biased significantly from the true values (green line) and falsely indicates a decrease in recombination rates the past, whereas the correction leads to a much more reasonable estimate (red line) which is only slightly underestimating the recombination rate. 500 ARGs of 8 haplotypes are sampled from *ARGweaver* and divided into 5 groups, and each estimate is obtained as the average over 100 ARGs from each group. The error bars represent 1 SE among the five estimates.

The bias mainly comes from the fact that type 1 recombinations do not result in tree change at all and are unreported by *ARGweaver*, which instead reports recombination events according to (2). As the probability of a type 1 event depends on the number of lineages, this induces the appearance of temporal recombination rate changes. The way to correct this is to introduce “effective counts” instead of naive counts of the recombination events in each time interval. For a recombination happening on branch *b* at time *t* on tree 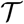, instead of counting it as one, we assign it the following weights:

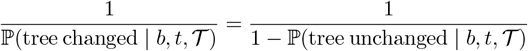

After applying this correction, we observe that the estimate are significantly closer to the true values (Figure 6) and no false evidence of temporal changes in the recombination rate are observed. In real data analyses it is, therefore, possible to infer changes in recombination rates using a combination of *ARGweaver* inferred trees and the correction derived here. We note that we did not test this procedure on simulated data with time-varying recombination rates because there is no available program to efficiently do such simulations.

## 4 Discussion

In this paper, we derived analytical formulae for the distribution of waiting distance to the next tree change and close approximations for the waiting distance to the next topology change under the SMC’ model. We use these results to show that tree transition omission in ARG/genealogy inference methods is a common problem, causing biases away from the SMC/SMC’ assumption that the waiting distance is exponentially distributed with intensity equal to the product of recombination rate and tree length. This challenges the use of such methods for making inferences about recombination rates and recombination rate evolution. The waiting distance between adjacent trees reported by *ARGweaver* is close to what is predicted by theory, and matches that expected for a valid Bayesian ARG sample. However, neither *Relate* nor *tsinfer* provide easily interpretable waiting distances between trees, and under realistic setting in humans, undercount topology changes and overestimate waiting distances.

We also highlight the use of ARG inference in understanding the temporal variation of recombination rates, by estimating the time-specific recombination rates using the correction proposed here, which can serve as a tool for understanding the evolution of recombination rates. However, we note that the constant population size demography model may not apply to most scenarios for real data, which will affect the calculation of the probability in (2). We argue that this problem can still be solved by re-calculating the probability given a demography model, and the extension is straightforward when the demographic model only involves a single population with changing size. Also, ARGs could be sampled using *ARGweaver-D* (Hubisz et al., 2020) instead of *ARGweaver* under a non-standard demography model. Finally, we note that importance sampling aproaches could be used to adjust for model misspecifications.

## Acknowledgments

We thank Aaron Stern and Debora Brandt for helpful discussions and Shang Liu for suggestions in programming. This research is supported in part by an NIH grant R35-GM134922 to YSS and NIH grant R01GM138634 to RN. YSS is a Chan Zuckerberg Biohub Investigator.

## Appendices

### A Proofs

#### A.1 Proof of Preposition 1 and Preposition 2

*Proof*. To simplify the notation we define *i* = *A_n_*(*t*) so that *t* ∈ [*σ*_*i*+1_,*σ_i_*], which gives us the following decomposition of the integral:

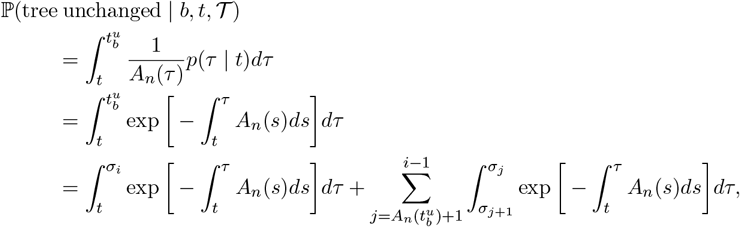

where *p*(*τ* | *t*) stands for the probability distribution of the rejoining time *τ* given the recombination time *t*. We can simplify the first term and summands of the second term as follows:

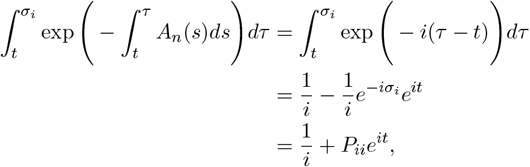

and

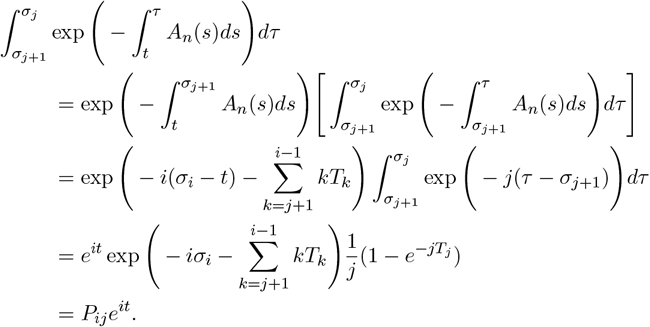

*Proof.* The probability of type 1 event on a branch *b*, which is the probability that a recombination not to change the tree conditioning on it happening on branch *b*, can be found by integrating over the breakpoint time, *t*, with respect to its conditionally uniform distribution on the branch.

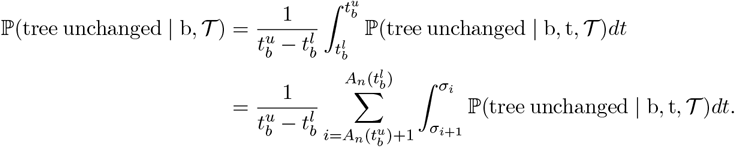

As for the summands, we apply our previous result to get:

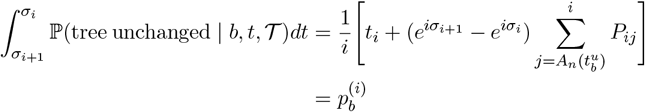

#### A.2 The proof of Theorem 3

For each branch *b* in the tree 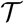 we call the lineage which coalesces with it *b*’. And we denote the parental lineage of *b* and *b*’ by *c*.

*Proof.* We note that the steps are basically the same with the proof of Theorem 1, which is to approach 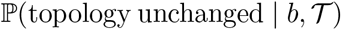 by integrating 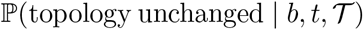, and then use the law of total probability to get 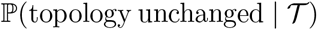.

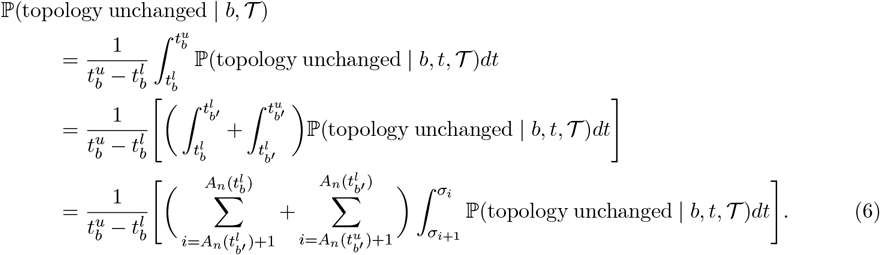

We need to break cases by whether [*σ*_*i*+1_,*σ_i_*] is a time period which is shared by both branch *b* and *b*’ or not.

**First case**: 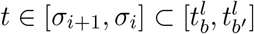,:

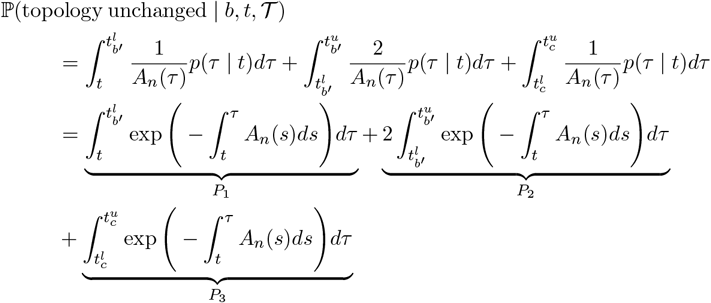

and we can simply the summands as follows:

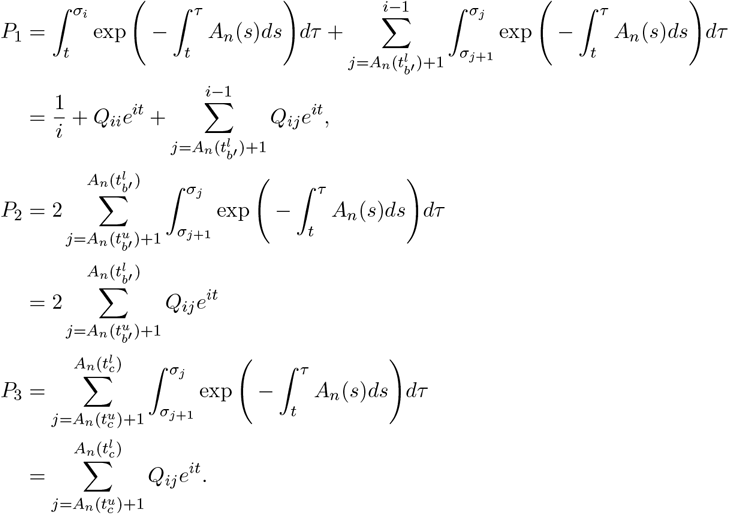

Plugging these terms back in, we obtain

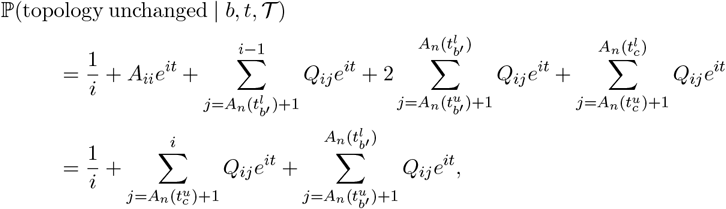

and

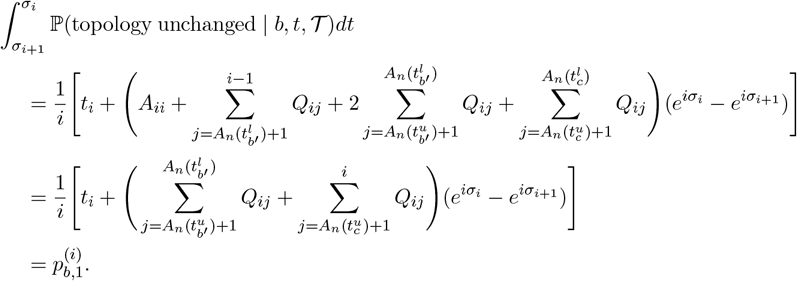

**Second case:** 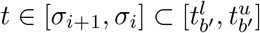

By similar arguments as above, we obtain

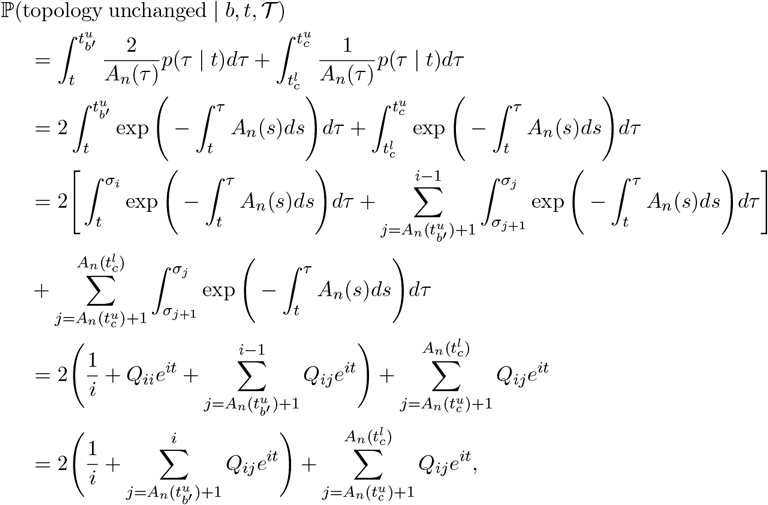

and

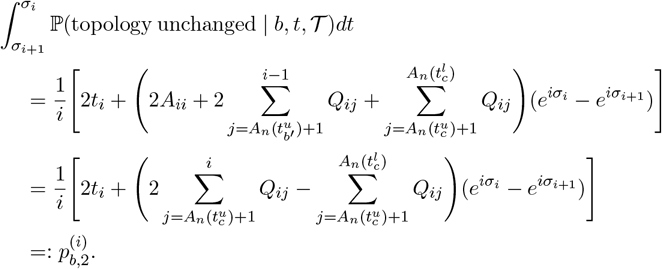

Now, we can plug these results into (6) to get

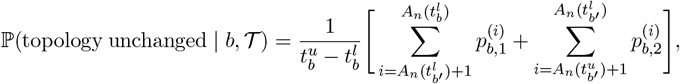

which leads to,

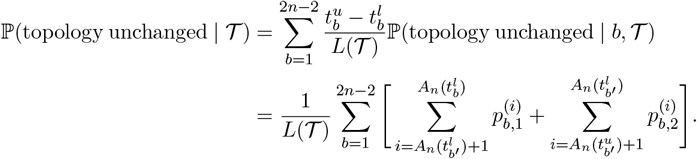

